# BTK operates a phospho-tyrosine switch to regulate NLRP3 inflammasome activity

**DOI:** 10.1101/864702

**Authors:** Zsófia A. Bittner, Xiao Liu, Sangeetha Shankar, Ana Tapia-Abellán, Hubert Kalbacher, Liudmila Andreeva, Matthew Mangan, Peter Düwell, Marta Lovotti, Karlotta Bosch, Sabine Dickhöfer, Ana Marcu, Stefan Stevanović, Franziska Herster, Markus W. Löffler, Olaf-Oliver Wolz, Nadine A. Schilling, Jasmin Kümmerle-Deschner, Samuel Wagner, Anita Delor, Bodo Grimbacher, Hao Wu, Eicke Latz, Alexander N. R. Weber

## Abstract

Activity of the NLRP3 inflammasome, a critical mediator of inflammation (*1*), is controlled by accessory proteins *(2, 3)*, post-translational modifications *(4, 5)*, cellular localization *(6, 7)* and oligomerization *(8)*. How these factors relate, is unclear. We show that the established drug target, Bruton’s Tyrosine Kinase (BTK) *(2, 9)*, integrates several levels of NLRP3 regulation: BTK phosphorylation of four conserved tyrosine residues, by neutralizing the charge of a polybasic linker region, weakens the interaction of NLRP3 with Golgi phospholipids and may thus guide NLRP3 cytosolic localization. BTK activity also promotes NLRP3 oligomerization and subsequent formation of inflammasomes. As NLRP3 tyrosine modification ultimately also impacts on IL-1β release, we propose BTK-mediated, charge-switch-based NLRP3 regulation as a novel and therapeutically tractable step in the control of inflammation.

**One Sentence Summary:** Multi-phosphorylation of NLRP3 by Bruton’s tyrosine kinase modulates NLRP3 cellular localization, inflammasome assembly, and IL-1β release.

## Introduction

Inflammation mediated via the NLRP3 inflammasome supports the resolution of infections and sterile insults but also contributes to pathology in multiple human diseases such as Cryopyrin-associated periodic fever syndromes (CAPS), gout, stroke or Alzheimer’s disease, and atherosclerosis *(10, 11)*. Thus, the activity of the NLRP3 inflammasome, a powerful molecular machine maturing IL-1 family cytokines via the activity of caspase-1, is tightly controlled at several levels: At the structural level, recent cryo-EM studies demonstrated that the 3D conformation of NLRP3 is critical for NLRP3 oligomerization and may depend on ADP/ATP binding *(8)*. In addition, NLRP3 binding proteins, such as NEK7, have been shown to have an impact on inflammasome activity *(3, 12)*. Moreover, post-translational modifications of NLRP3 enhance or reduce its activity by only partially elucidated mechanisms *(5)*. Finally, NLRP3 interacts dynamically with subcellular organelles such as the trans-Golgi network (TGN): On the one hand, a polybasic region in the linker between NLRP3 pyrin (PYD) and NACHT domains controls tethering of NLRP3 to negatively charged phosphatidylinositol-4-phosphate (PtdIns4P) at the disperse TGN *(6)*; on the other hand, dissociation from the TGN into the cytosol was proposed a requirement for the nucleation of larger NLRP3 oligomers and, subsequently, the assembly of complete inflammasome complexes, which include the adaptor ASC and IL-1β maturation enzyme, caspase-1 *(7)*. The cues instigating this shift in localization are not well understood. Generally, how these multiple layers of NLRP3 regulation are related or even integrated at the cellular as well as at the molecular level remains unclear. If individual regulators were to provide this integration, they could be valuable targets to modulate inflammasome activity.

We and others have recently identified Bruton’s tyrosine kinase (BTK) as a novel and therapeutically relevant NLRP3 regulator *(2, 13)*, which is rapidly activated upon NLRP3 inflammasome stimulation, and interacts with NLRP3 and ASC in overexpression systems. Its genetic ablation led to reduced IL-1β secretion *in vitro* and, importantly, in human patients *ex vivo (2)*. BTK is a well-known cancer target for which FDA-approved inhibitors such as ibrutinib exist (reviewed in Ref. *(9)*). Based on the molecular mechanisms and chronic inflammatory processes observed in the pathology of many cancers, targeting NLRP3 via BTK also appears as an attractive therapeutic option in other diseases.

Here we report that BTK directly modifies four NLRP3 tyrosine (Y) residues in the PYD-NACHT polybasic linker, neutralizing positive charge. This charge-switch weakens phospholipid interactions and prompts dissociation from Golgi membranes. Consequently, ablation of BTK kinase activity or tyrosine mutation decreased the formation of cytosolic NLRP3 oligomers, complexes with ASC and IL-1β release, respectively. Our data suggest that this BTK-mediated tyrosine phosphorylation affects the activity of the NLRP3 inflammasome by modifying linker charge, subcellular localization, inflammasome assembly and ultimately IL-1β secretion. BTK thus emerges as an important regulation hub for the activation of the NLRP3 inflammasome at multiple levels, which could be simultaneously targeted via BTK kinase inhibitors.

## Results

### BTK deficiency coincides with reduced NLRP3 tyrosine phosphorylation

Based on previous work (2), we hypothesized that BTK and NLRP3 may engage in a direct kinase-substrate relationship, whose elucidation might unravel novel molecular aspects of NLRP3 inflammasome regulation. We sought to test this in Btk-deficient primary murine BMDMs and in PBMCs from patients with the genetic BTK deficiency, X-linked agammaglobulinemia (XLA). As expected, IL-1β release upon nigericin stimulation was significantly reduced in BTK-deficient BMDMs and patient-derived PBMC, respectively (Fig. 1A, B). Interestingly, in BMDMs, endogenous NLRP3 precipitated and interacted with endogenous BTK in WT but not Btk or Nlrp3 KO BMDM (Fig. 1C), irrespective of the BTK kinase inhibitor ibrutinib. Similarly, BTK co-immunoprecipitated with NLRP3 in PBMCs from healthy donors (HDs) (Fig. 1D). Thus, in both human and murine primary immune cells BTK and NLRP3 interact independently of nigericin stimulation. This was also confirmed with a cell-free *in vitro* pull-down of recombinant purified NLRP3 and BTK proteins (Fig. 1E, Ref. (8) and Methods). We next tested whether BTK is able to phosphorylate NLRP3 upon nigericin treatment. In murine BMDMs (Fig. 1F) immunoprecipitated NLRP3 became rapidly phospho-tyrosine (p-Y)-positive, in cells expressing Btk but not in *Btk* or *Nlrp3* KO cells. Similarly, NLRP3 phosphorylation was also observed in healthy donor PBMCs (Fig. 1G), and lower in XLA PBMCs (Fig. S1A). Importantly, treatment with λ-phosphatase abolished p-Y reactivity, further confirming the phospho-antibody specificity. Thus, BTK and NLRP3 interact endogenously in primary immune cells and BTK promotes NLRP3 tyrosine-phosphorylation upon nigericin stimulation.

**Figure 1:**
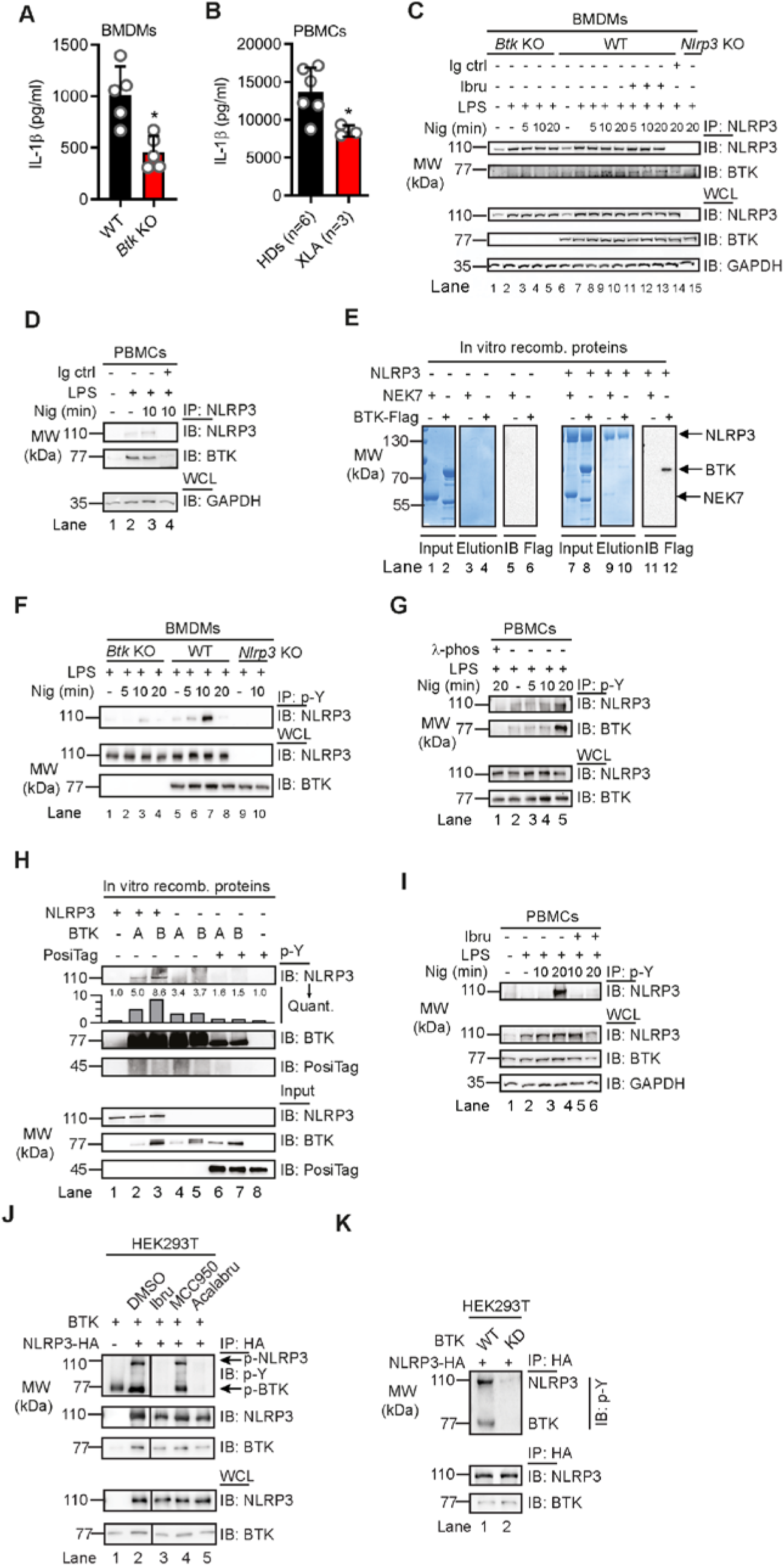
NLRP3 directly interacts and is tyrosine phosphorylated by BTK. (A, B) IL-1β release (triplicate ELISA) from WT vs Btk KO BMDMs (A, n=5 each) or XLA vs healthy donor (HD) PBMCs (B, n=3-6). (C, D) Co-IP of NLRP3 from WT, Btk KO or Nlrp3 KO BMDM (n=3) or ibrutinib-treated PBMC lysates (n=2). (E) *In vitro* pulldown of FLAG-tagged BTK or His-SUMO-tagged NEK7 by MBP-tagged NLRP3 (n=3). (F, G) as in C and D, respectively, but with anti-phospho-tyrosine (p-Y) IP (n=3 or 5, respectively). (H) as in E but using two different commercial suppliers, A and B, of recombinant BTK. PosiTag = specificity control. (I) as in G but with ibrutinib pre-treatment (n=2). (J, K) IPs from HEK293T cells transfected with NLRP3 and BTK WT or kinase dead (KD) constructs, or treated with inhibitors (n=2 each). A and B represent combined data (mean+SD) from ‘n’ biological replicates (each dot represents one mouse or patient/HD). C-K are representative of ‘n’ biological (HD or mouse) or technical replicates. * p<0.05 using Student’s *t*-test (A) or one-way ANOVA with Dunnett’s correction (B).

### BTK kinase activity is required for NLRP3 tyrosine phosphorylation

We next tested whether BTK kinase activity was required for NLRP3 tyrosine phosphorylation. Two independent cell-free *in vitro* setups showed that the presence of BTK was necessary and sufficient for NLRP3 p-Y modification (Figs. 1H and S1B). However, in both PBMCs and the *in vitro* setup NLRP3 tyrosine phosphorylation was blocked by ibrutinib treatment (Figs. 1I and S1B) and, thus, dependent on BTK kinase activity. Next, HEK293T cells were transfected with NLRP3 and BTK, and treated with or without BTK inhibitors. NLRP3 and BTK interacted independently of BTK kinase inhibitors, ibrutinib and acalabrutinib; however, NLRP3 tyrosine phosphorylation was abrogated in the presence of both BTK kinase inhibitors (Fig. 1J), consistent with results in primary BMDMs (*cf.* Fig. 1C). In contrast, the NLRP3-specific inhibitor, MCC950 (14), failed to prevent BTK-specific interaction and NLRP3 p-Y positivity (Fig. 1J). Interestingly, the expression of kinase-dead (KD) BTK (K430E mutation, see Ref. (9)) was not able to induce NLRP3 tyrosine phosphorylation, despite an intact interaction (Fig. 1K). Similar results were obtained for the *in vitro* cell-free setup (*cf.* Fig. S1B). Thus BTK kinase activity appears essential and sufficient for NLRP3 p-Y modification using primary immune cells, the HEK293T system or purified recombinant proteins, indicative of a direct kinase-substrate relationship.

### BTK phosphorylates four conserved tyrosine residues in the NLRP3 PYD-NACHT linker domain

To map the modified tyrosine residues in NLPR3, we compared Flag-tagged full-length with truncated NLRP3 constructs (15) of only the PYD (1-93), the extended NACHT domain (94-534), which includes an N-terminal linker domain (94-219) (8), and the LRR domain (535-1,036), see Figs. 2A and S2. BTK exclusively phosphorylated the NLRP3 extended NACHT construct (Fig. 2B, C), ruling out Y861 (5) as the phospho-site. Mutation of the nine tyrosines in the core NACHT domain (220-534), see Fig. 2D) to phenylalanine (F) did not impact the level of phospho-NLRP3 detected upon BTK co-expression (Fig. S3A, quantified in S3B). However, when the linker (94-219) tyrosines were targeted (Fig. 2E), mutated Y168 showed partial but significant reduction of the p-Y signal, as shown by conventional immunoblotting (Fig. 2F, quantified in G) and WES capillary electrophoresis analysis (Fig. 2H, quantified in S3C), respectively. Thus, Y168 emerged as a novel putative phospho-tyrosine site in NLRP3, specifically modified by BTK. Unfortunately, the linker region is not accessible to mass spectrometric analysis (data from (16) plotted in Fig. S3D). Therefore, to assess the phosphorylation of Y168 by alternative means, 15-mer peptides covering all linker tyrosines (*cf.* Fig 2E and Table S1) were incubated with His-tagged BTK to assess peptide phosphorylation in a cell-free system. Following BTK removal by anti-His beads, tyrosine phosphorylation of the peptides was visualized by dot blot analysis. While the majority of Y-containing peptides and all F-containing corresponding peptides were not phosphorylated (Fig. S3E), the Y168-containing peptide showed strong tyrosine phosphorylation (Fig. 2I). Of note, peptides containing Y136, Y140, or Y143 – either in combination or as single tyrosines – were also phosphorylated (Fig. 2I), similar to peptides containing the sequences in mouse NLRP3 (Y132, Y136, Y145 and Y164, see Fig. S3F). In HEK293T cells, combined mutations of Y136, Y140, Y143 (“3Y>F”) and additionally Y168 to phenylalanine (“4Y>F”) consequently led to a strong reduction of the p-Y signal, both in a FL-NLRP3 construct and when the linker was fused to an mCitrine yellow fluorescent protein-HA sequence (here termed ‘hLinker-Cit-HA’, Fig. 2J). Consistent with the only partial effect of a single Y168F mutation (*cf.* Fig. 2F-H), BTK appears to be able to specifically modify not only one, but at least four tyrosines – Y136, Y140, Y143 and Y168 – in the PYD-NACHT linker of human (and murine) NLRP3 *in vitro*. To gain structural insights, we mapped the sites in a recent cryo-EM structure of an NLRP3-NEK7 complex (Ref. (8), Fig. 2K and S4A). Interestingly, Y136, Y140 and Y143 were found in a helical region proposed to contact the PYD for ASC recruitment (8). Further, Y168, which maps to the vicinity of several likely pathogenic CAPS mutations (*cf.* Fig. S2), was adjacent to a putative ADP molecule and might thus influence nucleotide binding (Fig. 2L, M). Interestingly, all of the BTK-modified Y residues were strongly conserved in other NLRP3 sequences, further highlighting their potential functional relevance (Fig. S4B). Collectively, our results suggest that BTK directly modifies multiple highly conserved tyrosines within a functionally important domain in NLRP3 that could impact downstream steps in the assembly of the oligomeric inflammasome complexes.

**Figure 2:**
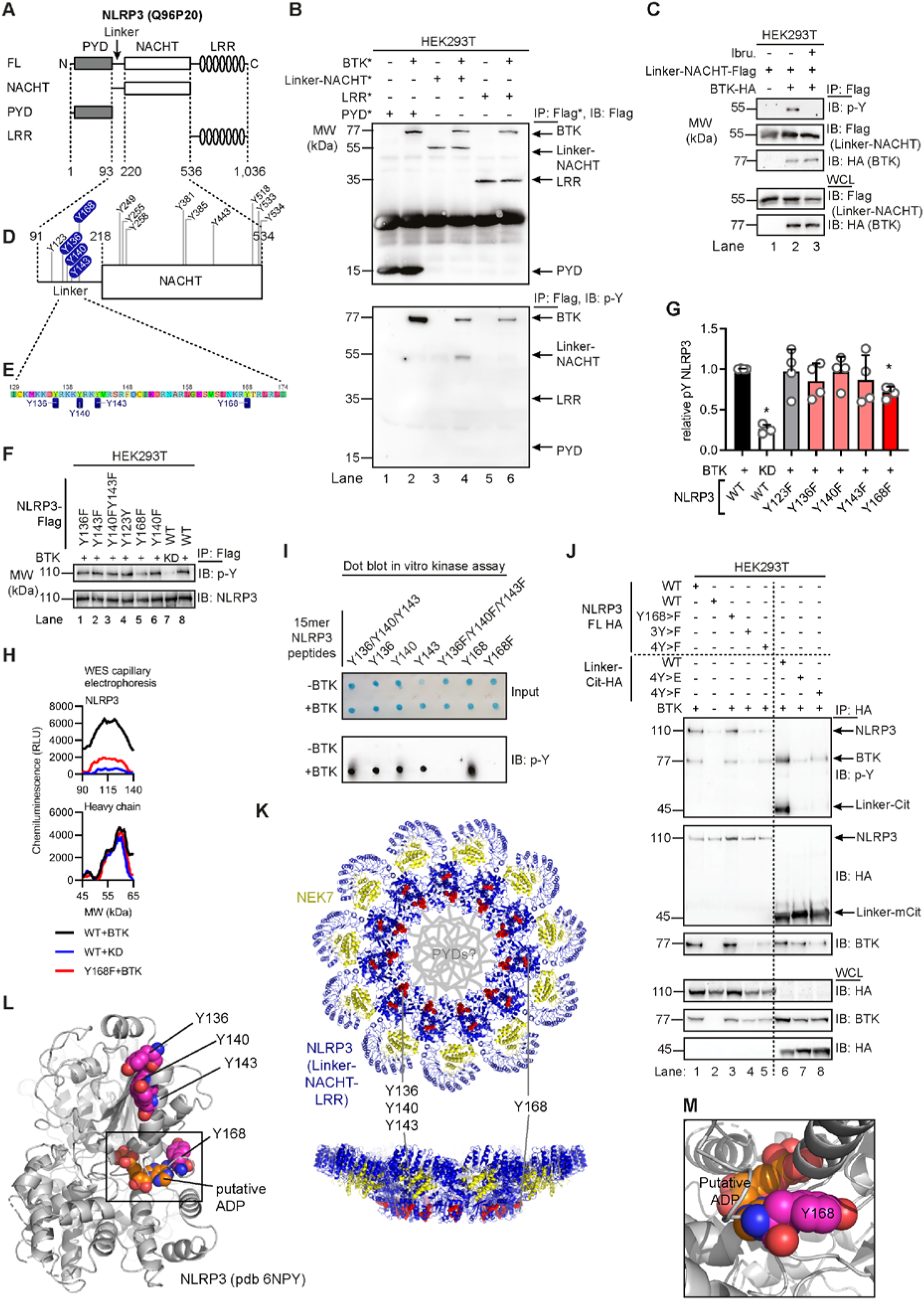
BTK phosphorylates the PYD-NACHT linker. (A) NLRP3 domains (UniProt ID Q96P20). (B) IP from HEK293T cells transfected with NLRP3 and BTK constructs (n=3). (C) as in B but including ibrutinib (n=3). (D) Positions of targeted tyrosine residues. (E) Linker region including polybasic motif. (F) as in B but using Y to F point mutants and WT or kinase-dead (KD) BTK plasmids (n=4). (G) Quantification of F. (H) WES capillary electrophoresis of IP p-NLRP3 (n=3). (I) Dot blot of BTK kinase assay with 15-mer synthetic peptides (n=3). (J) as in F but also NLRP3 linker (WT or Y-mutated) fused to mCitrine-HA (n=3). (K) Tyrosines (red) highlighted in model of NLRP3 (blue)-NEK7 (yellow) complex (pdb: 6NPY). Close up view on dimer interface (L) and putative nucleotide binding site (M). G represents combined data (mean+SD) from ‘n’ biological replicates (each dot represents one replicate). B, C, F, H-J are representative of ‘n’ technical replicates. * p<0.05 according to one sample *t*-test (G).

### BTK-mediated phosphorylation sites affect PI4P binding

Chen and Chen recently identified a ‘polybasic’ motif (K127-130 in mouse; K127 and K129-130 in human NLRP3) as critical for NLRP3 charge-dependent binding to Golgi phosphatidylinositol-4-phosphates (PI4Ps) and inflammasome nucleation (6). Unexpectedly, Y136, Y140 and Y143 precisely mapped to this polybasic motif in the NLRP3 PYD-NACHT linker, and interdigitate with the basic residues mediating the proposed NLRP3 interaction with negatively charged membrane phospholipids (Fig. 3A). The mutation of three positively charged residues (K127, K129, K130) in the mNLRP3 polybasic region to alanine (K>A) is sufficient to abrogate PI4P binding (6). We therefore hypothesized that BTK phosphorylation of Y136, Y140 and Y143 might weaken the proposed charge attraction of NLRP3 with Golgi PI4Ps (6, 7). Charge computations suggested that at cytoplasmic pH 7.4, the net charge of the NLRP3 polybasic linker sequence shifts from +7.28 (unphosphorylated) to +2.01 when Y136, Y140 and Y143 are phosphorylated in human NLRP3 (Fig. 3B), and from +8.33 (unphosphorylated) to +3.06 (3x phosphorylated) in mouse NLRP3. To confirm this charge shift, synthetic phospho- and non-phospho versions of the Y136/Y140/Y143-containing peptides from both human and mouse NLRP3 were studied in pH titrations, demonstrating the phospho-Y-peptide to be significantly more acidic/negatively charged (Fig. 3C). Furthermore, in the NLRP3-NEK7 structure the 3x phospho-modification is calculated to cause a significant change in the computed surface charge of a ‘hypothetically active’ NLRP3 state (8) towards negative values (Fig. S5A). We therefore hypothesized tyrosine phosphorylation by BTK might weaken interactions with the negatively charged PI4Ps. Consistently, the binding of 4xY>E, i.e. phospho-mimetic, mutant human NLPR3 linker-Cit-HA fusion protein to PI4P beads (6) was reduced compared to the corresponding WT construct (Fig. 3D). This was further confirmed by the fusion of the murine NLRP3 polybasic region to GFP-Flag (mPBR-GFP-Flag, construct as in Ref. (6)): Despite equal expression, the Y>E construct bound PI4P beads less strongly than WT (Fig. 3E). Reduced binding was also observed for the corresponding K>A mutant, which is known to be defective in PI4P binding based on charge neutralization (6). To test whether BTK activity might also affect Golgi localization, we fractionated different stimulated BMDM lysates into S100 (soluble, cytosol), P5 (heavy membrane, Golgi and mitochondria) and P100 (light membrane, ER and polysomes) fractions and probed them for NLRP3. BTK, whose PH domain also binds PIPs (references in (17), was also analyzed and found to localize similarly to NLRP3 (Fig. 3F, Fig. S5B), in keeping with earlier interaction analyses (*cf.* Fig. 1C, F). As expected (6), in both WT and Btk KO BMDMs, NLRP3 was detectable in the P5 fraction upon LPS treatment, indicating that BTK is not essential for initial NLRP3 localization towards P5 membranes (Fig. 3F, Fig. S5B). However, nigericin stimulation in WT BMDMs coincided with progressive depletion of NLRP3 from the P5 fraction within 20 min, consistent with the timing of phosphorylation in these cells (*cf.* Fig 1F). Conversely, Btk KO BMDMs did not show dissociation within this time frame, suggesting the NLRP3 interaction with Golgi membranes remains more stable in the absence of BTK, possibly due to intact charge-mediated PI4P-interactions (6). Collectively, these experiments show that both, the BTK-modified tyrosine positions in the NLRP3 linker and the presence of BTK, have an impact on NLRP3 linker-PI4P binding and subcellular fractionation of NLRP3.

**Figure 3:**
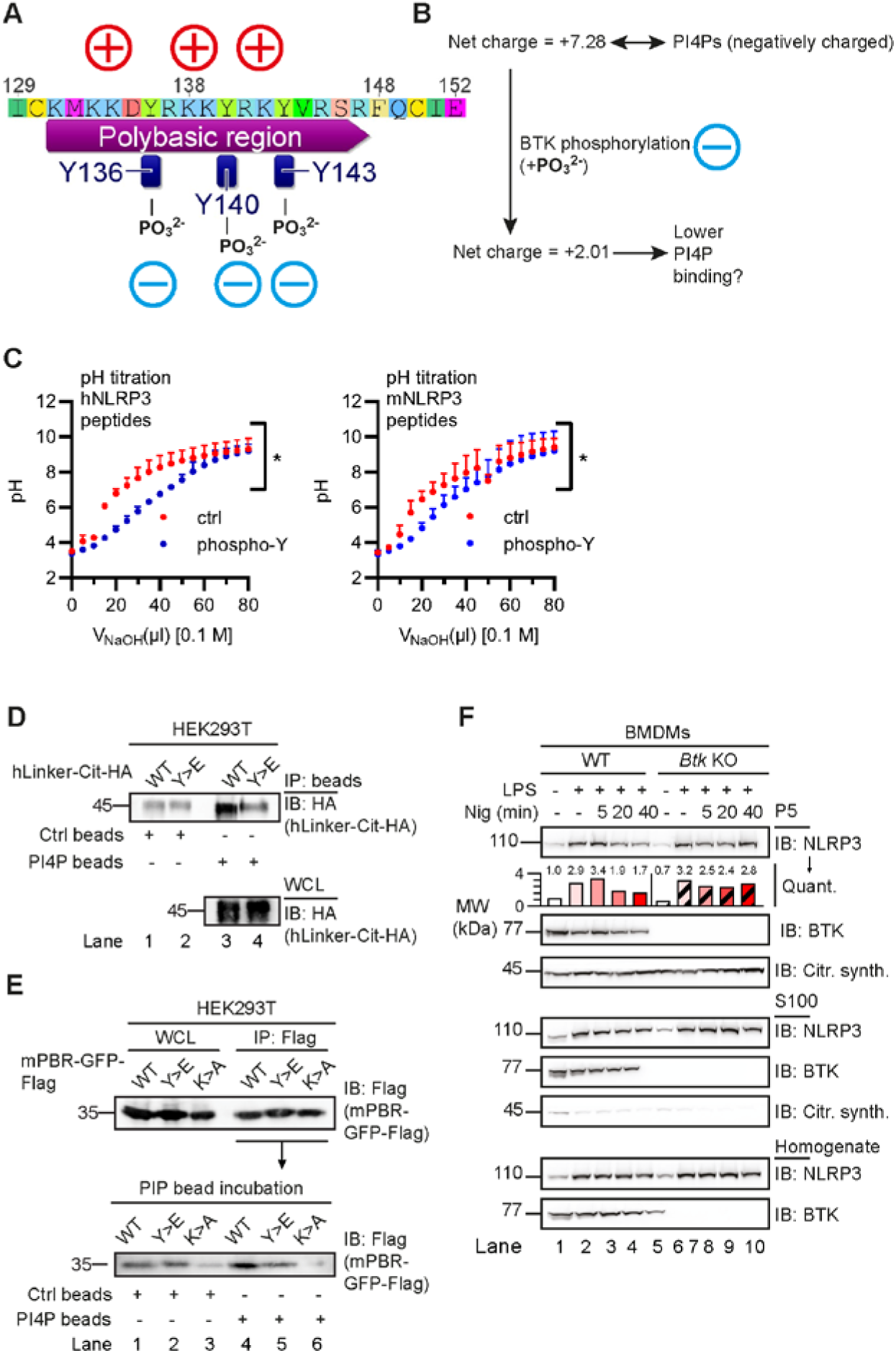
BTK phosphorylation of the NLRP3 polybasic motif enables Golgi/PI4P dissociation. (A, B) Charge distribution (A) and ProtPi net charge computation (B) of unmodified and 3x phospho-peptide polybasic human NLRP3 linker. (C) pH titration of peptides encompassing the polybasic motifs of human or murine NLRP3 as phospho- (blue) or non-phosphorylated control (red) peptide (n=3). (D) NLRP3 linker-Cit-HA constructs precipitated with PI4P beads (n=3). (E) as in D but murine NLRP3 polybasic region fused to GFP-HA (mPBR-GFP-HA, n=3). (F) Subcellular fractionation of nigericin-treated WT or Btk KO BMDM lysates (n=3). C represents combined data (mean+SD) from ‘n’ biological replicates. In D-E one representative example of ‘n’ technical replicates is shown. * p<0.05 according to two-way ANOVA (C).

### BTK kinase activity affects NLRP3 oligomerization, ASC interaction and IL-1β release

A recent study proposed that NLRP3 release from the Golgi was required for ASC engagement and full inflammasome assembly (7). As Golgi depletion was lower in the absence of BTK (*cf.* Fig. 3F), we explored whether BTK kinase activity also affects subsequent full inflammasome assembly, e.g. by recruiting ASC into complex formation. Indeed, native PAGE of nigericin-stimulated WT and Btk KO BMDM lysates (Fig. 4A) or BTK inhibitor- vs vehicle-treated Pycard (ASC) KO (Fig. 4B) showed reduced NLRP3 oligomers in BTK-deficient or –inhibited samples, and lower ASC cross-linking (Fig. 4C). Size exclusion chromatography of untreated WT cell lysates showed that BTK and NLRP3 co-eluted in the high MW fraction (>1,100 KDa). Consistent with native PAGE in Btk KO or WT lysates from ibrutinib-treated cells, elution shifted to lower molecular weight complexes (Fig. 4D, E). Ablation of BTK activity, thus, appeared to reduce the subsequent ability of NLRP3 to oligomerize into large MW cytosolic inflammasomes and to assemble with ASC. To show that BTK-modified tyrosine residues – and the effects of BTK described so far – also had an impact on IL-1β release, NLRP3-deficient immortalized macrophages were retrovirally transduced with WT or tyrosine-mutated (4xY>F) NLRP3-T2A-mCherry constructs, allowing for cell sorting for equal protein expression (Fig. 4F). Compared to WT-transduced cells, cells with the 4xY>F construct failed to restore nigericin- and R837 (imiquimod)-dependent IL-1β release (Fig. 4G). Conversely, TNF release, which is NLRP3-independent (2), was comparable between cell lines (Fig. 4H). Likewise, IL-1β release upon stimulation with the NLRP3- and BTK-independent (2) AIM2 inflammasome stimulus, poly(dA:dT), was comparable between WT and mutant cells (Fig. 4G, H). The data show that the BTK-modified tyrosine positions identified here play an important role for full NLRP3 inflammasome activity and subsequent IL-1β release.

**Figure 4:**
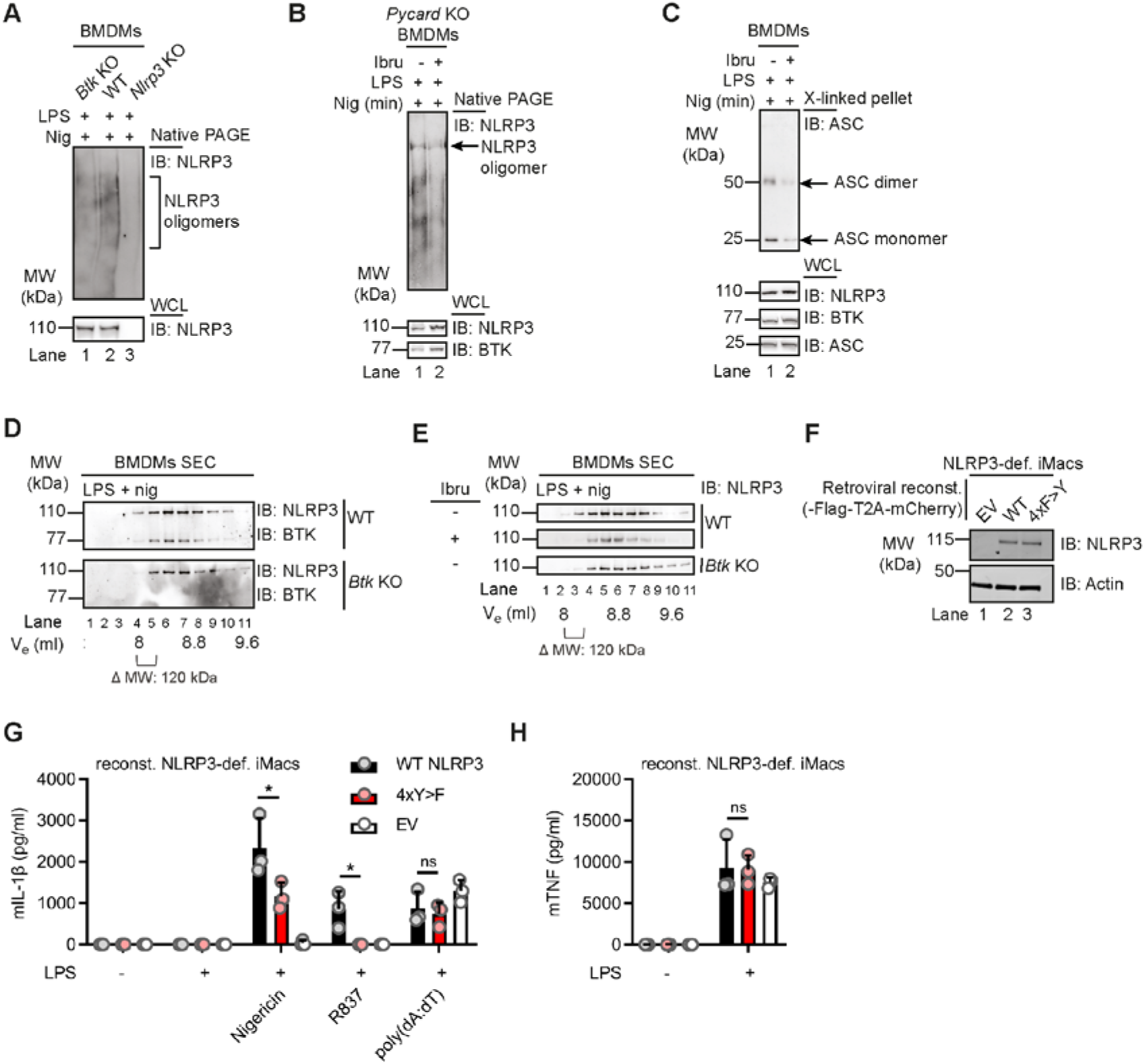
BTK modification affects NLRP3 oligomerization and IL-1β release. (A-C) WT, Btk KO, Nlrp3 KO or Pycard (ASC) KO BMDMs stimulated and respective lysates analyzed directly by native PAGE (A, n=2 and B, n=4) or ASC cross-linked in the pellet (C, n=4) and/or analyzed by SDS-PAGE. (D) as in A but size exclusion chromatography (SEC) fractions (n=3). (E) as in D comparing inhibitor treated WT BMDM or Btk KO BMDM lysates (n=3). (F-H) NLRP3 expression levels (F), IL-1β (G) or TNF (H) release from WT or 4xY>F NLRP3-reconstituted NLRP3-deficient iMacs (n=3). G-H represent combined data (mean+SD) from ‘n’ technical replicates. A-F are representative of ‘n’ biological (mice) or technical replicates. * p<0.05 according to one-way ANOVA (H) or ANOVA with Sidak correction (G).

## Discussion

The mechanism of activation of the NLRP3 inflammasome has been intensely studied for some time and recent work has unraveled a critical role of the dynamic localization of NLRP3 at the Golgi in the regulation of its activity *(6),(7)*. Using both biochemical and cellular assays in human and murine primary cells, we found that BTK directly participated in these processes at the level of NLRP3, providing novel molecular insights: (i) direct phosphorylation of four conserved and functionally important tyrosine residues in the NLRP3 polybasic linker motif (ii) affected linker charge; (iii) phospho-mimetic mutation weakened NLRP3-PI4P interactions, presumably by neutralization of the linker net surface charge. In line with this, (v) in the presence of BTK, NLRP3 retention to the Golgi is seemingly less, whereas (vi) NLRP3 inflammasome oligomerization, ASC association, and IL-1β secretion are higher. Finally, (vi) mutation to the non-phosphorylatable phenylalanine abrogated full IL-1β release, highlighting the importance of these tyrosines. Our data suggest that BTK-mediated phosphorylation of multiple NLRP3 tyrosines may thus serve as a kind of molecular switch, tuning NLRP3 charge and subsequently localization and inflammasome assembly. Modification, localization, and oligomerization of NLRP3 have been recognized to be important. However, they were supposed to be hierarchically separate layers of NLRP3 inflammasome regulation and, hence, of inflammation. Our data indicate that some of these layers may be integrated and interconnected by BTK: By decoding the most basal determinants such as protein sequence, BTK appears to integrate post-translational modifications, surface charge, interaction with organelles, and ultimately assembly of a highly oligomeric molecular machinery (Fig. S6). Such a dense and interconnected network of regulation would be in line with the critical need to tightly control excessive IL-1β release to prevent pathologies. Of course, other proteins may also participate in this process and we cannot formally rule out effects of BTK at the level of ASC. Additionally, the events upstream of BTK await elucidation outside the scope of this present work. A recent report described a reversal of the role of BTK in inflammasome activation by unphysiologically high LPS priming, see supplemental discussion. Our data clearly indicate that BTK is directly involved in licensing full IL-1β release via the regulatory events characterized here. Furthermore, NLRP3 phosphorylation may represent a biomarker, and possibly a therapeutic target, for early NLRP3 activation. Collectively, our work provides both a rationale as well as manifest targeting strategies that may be applied to block excess IL-1β production in acute inflammasome-related diseases.

## Abbreviations

AIM2: Interferon-inducible protein absent in melanoma 2
ASC: Apoptosis-associated speck-like protein containing a Caspase activation and recruitment domain (CARD)
BMDM: bone marrow-derived macrophages
BTK: Bruton’s Tyrosine Kinase
F: Phenylalanine
FDA: Food and Drug Administration
CAPS: Cryopyrin-associated periodic syndrome
GM-CSF: Granulocyte-macrophage colony-stimulating factor
HD: healthy (blood) donor
HEK: human embryonic kidney
IFN: Interferon
IL: Interleukin
IP: immunoprecipitation
KD: kinase-dead
LPS: Lipopolysaccharide
LRR: leucine-rich repeat
NEK7: NIMA related kinase 7
NACHT: NAIP, CIITA, HET-E and TEP1
NLR: Nod-like receptor
NLRP3: NACHT, LRR and PYD domains-containing protein 3
PBMC: peripheral blood mononuclear cell
PH: Pleckstrin homology
PI4P: phosphatidylinositol-4-phosphate
PMA: Phorbol-12-myristate-13-acetate
p-Y: phospho-tyrosine
PYD: Pyrin domain
SH: Src homology
TGN: trans-Golgi network
TH: Tec homology
TLR: Toll-like receptor
TNF: Tumor necrosis factor
XLA: X-linked agammaglobulinemia
Y: Tyrosine.

## Acknowledgments

We gratefully acknowledge Ulrich Wulle for help with peptide synthesis and Yamel Cardona Gloria for helpful comments. We thank Xiaowu Zhang and Felix Meissner for helpful advice on kinase target residue identification and mass spectrometry, respectively. We thank all study subjects and their families for participating in the study.

## Author contributions

ZAB, XL, SD, HK, KB, LA, AM, MM, PD, ML, FH, SS, ATA, OOW, NAS, SW and ANRW performed experiments; ZAB, XL, SD, HK, LA, SS, ML, MM, PD, FH, NAS, SW and ANRW analyzed data; MWL, JKD, AD and BG were involved in patient recruitment and sample acquisition; ZAB and ANRW wrote the manuscript and LA, PD, MWL, SS, ATA, SW, HW and EL provided valuable comments. All authors approved the final manuscript. ANRW and ZAB conceived and coordinated the study. S.D.G.

## Data and materials availability

All data is available in the main text or the supplementary materials. Materials are available upon reasonable request.

