## Supplemental information for "BTK operates a phospho-tyrosine switch to regulate NLRP3 inflammasome activity"

**This PDF file includes:**

Supplementary discussion
Materials and Methods
Supplementary Text
Figs. S1 to S6
Tables S1 to S2
References cited in supplement

### Supplementary discussion

A recent report by Mao et al suggested that BTK deficiency augments NLRP3 inflammasome activation and causes IL-1β-mediated colitis" (www.jci.org/articles/view/128322). As at first glance, these results appear to contradict our work, we discuss the relationship between this report and our present work here. The two first and independent reports on a role of BTK in NLRP3 inflammasome regulation (Ito, Shichita et al. 2015; Liu, Pichulik et al. 2017) clearly indicated the role of a positive regulator: We and Ito *et al.* both observed reduced IL-1β release in BTK-deficient mouse cells, whole animals and cell lines. Additionally, primary cells from BTK-deficient (X-linked agammaglobulinemia) and BTK-inhibitor-treated patients showed the same effect. Mao *et al.* instead propose that BTK acts conversely, as a negative NLRP3 regulator. This is based on their results from human and mouse BTK-deficient cells primed with 200 ng/ml LPS, which secreted more mature IL-1β than WT cells. Conversely, at LPS concentrations <100 ng/ml – which are more physiological (Zweigner, Gramm et al. 2001; Copeland, Warren et al. 2005) – , they observed lower IL-1β under BTK ablation, in actual agreement with our data. Mao et al. suggest that, in the absence of BTK, low LPS may insufficiently prime the inflammasome. However, under the LPS concentrations we used, *IL1B* mRNA induction, an important consequence of priming, and secretion of inflammasome-independent cytokines, were always comparable in WT and BTK-deficient murine or human samples (Liu, Pichulik et al. 2017). Additionally, in their work NLRP3 oligomerization and adaptor recruitment occur by LPS alone, i.e. without an NLRP3-specific stimulus, indicating that the excessive LPS stimulation used may blur the lines between TLR priming and NLRP3 activation. Mechanistically, Mao *et al.* go on to implicate BTK-mediated deactivation of the NLRP3-activating phosphatase PP2A under these conditions. It will be interesting to explore this novel mechanism in relationship to direct tyrosine phosphorylation of NLRP3 by BTK which we recently investigated (Bittner, Liu et al. 2019). Finally, an aggravated colitis phenotype observed by Mao *et al.* in BTK-deficient (and hence B cell-depleted) animals in vivo appears to argue for a negative inflammasome role and to parallel increased colitis in XLA and BTK inhibitor-treated leukemia patients. However, a genetically- or BTK inhibitor-induced lack of (regulatory) B cells is well known to exacerbate colitis in mice and humans (Yanaba, Yoshizaki et al. 2011; Wang, Ray et al. 2015; Kondo, Shaim et al. 2018) and thus a likely and critical confounder in their experiments. In our opinion even the presented in vivo evidence therefore does not unequivocally support the conclusion of BTK as an exclusively negative regulator of NLRP3 inflammasome activity. Nevertheless, this recent work points to a possible influence of TLR signaling in 'tuning' BTK for inflammasome regulation. Certainly, our and their studies warrant the further exploration of a possible 'rheostat role' of BTK that may tune NLRP3 activation.

### Materials and Methods

**Reagents.** Nigericin and Lipopolysaccharide (LPS) were purchased from Invivogen, ATP from Sigma, ibrutinib and acalabrutinib from Selleckchem, recombinant granulocyte-macrophage colony-stimulating factor (GM-CSF) from Prepro-Tech, Ficoll from Merck Millipore. Peptides (synthesized in house) and antibodies are listed in Tables S1 and S2, respectively.

**Peptides.** Synthetic peptides were produced by standard 9-fluorenylmethyloxycarbonyl/tert-butyl strategy using peptide synthesizers P11 (Activotec) or Liberty Blue (CEM Corporation). Purity was assessed by reversed phase HPLC (e2695, Waters, Eschborn, Germany) and identity affirmed by nano-UHPLC (UltiMate 3000 RSLCnano) coupled online to a hybrid mass spectrometer (LTQ Orbitrap XL, both Thermo Fisher). Lyophilized peptides were purified by standard HPLC. For certain peptides a pH titration with 0.1 M NaOH was performed using standard procedures. For *in vitro* assays peptides were dissolved at 10 mg/ml in dimethyl sulfoxide (DMSO) and diluted 1:10 in bidestilled $H_2O$. Frozen aliquots were further diluted in cell culture medium and sterile filtered if necessary.

**Plasmid constructs.** ASC, NLRP3 and BTK coding sequences in pENTR clones were generated as described in (Wang, El Maadidi et al. 2015). Truncated Flag-tagged NLRP3 constructs were a kind gift of F. Martinon, Lausanne, Switzerland (Mayor, Martinon et al. 2007). Constructs for the human PYD-NACHT linker (residues 94-219) fused to mCitrine-HA or the murine polybasic motif (residues 127-146) in the context of Flag-GFP (as in (Chen and Chen 2018)) were synthesized by GeneWiz. Point mutations in BTK and NLRP3 were subsequently introduced using QuikChange II Site-Directed Mutagenesis Kit from Agilent Technologies, following the manufacturer´s instructions. Presence of the desired mutation and absence of unwanted regions in the entire CDS was confirmed by automated DNA sequencing.

**Study subjects and blood sample acquisition.** CAPS patients were recruited at the Department of Pediatrics, University Hospital Tübingen and XLA patients at the Centre of Chronic Immunodeficiency, University Hospital Freiburg, healthy blood donors at the Interfaculty Institute of Cell Biology, Department of Immunology, University of Tübingen. All patients and healthy blood donors included in this study provided their written informed consent before study participation. Approval for use of their biomaterials was obtained by the respective local ethics committees, in accordance with the principles laid down in the Declaration of Helsinki as well as applicable laws and regulations. XLA patients were clinically identified and genetically characterized as described in (Liu, Pichulik et al. 2017).

**Mice.** *Btk* KO (Khan, Alt et al. 1995) and *NLRP3* KO (Jackson stock No: 021302) and wild type C57BL/6J (Jackson) colonies were maintained in specific-pathogen free conditions under regular hygiene monitoring. All animal experiments were approved by local authorities and performed in accordance with local institutional guidelines and animal protection laws, including specific locally approved protocols for sacrificing.

**Cell culture.** All cells were cultured at 37 °C and 5% $CO_2$ in DMEM or RPMI supplemented with 10% fetal calf serum, ʟ-glutamine (2 mM), penicillin (100 U/ml), streptomycin (100 µg/ml) (all from Thermo Fisher). They were free of mycoplasma contamination and monitored regularly using a PCR-based assay.

**Isolation and stimulation of primary human immune cells.** Peripheral blood mononuclear cells (PBMCs) from healthy donors and patients were isolated from whole blood using Ficoll density gradient purification, primed with 10 ng/ml LPS for 3 h, and in some cases treated with 60 µM ibrutinib for 15 min before stimulation with 15 µM nigericin for the indicated periods of time.

**Generation of primary bone marrow-derived macrophages (BMDMs) and NLRP3 stimulation.** Bone marrow (BM) cells were isolated from femurs and tibiae of 8-12 week old mice, grown and differentiated using GM-CSF (M1 polarization) as described (Liu, Pichulik et al. 2017). BMDMs were primed with 100 ng/ml LPS for 3 h and either treated with 60 µM ibrutinib for 15 min or stimulated directly with 5 µM nigericin for the indicated time-periods.

**Expression and purification of recombinant BTK, NEK7 and NLRP3.** The plasmids encoding NLRP3 with the deleted pyrin domain (amino acids 134–1034) for MBP-fusion protein expression in Bac-to-Bac system (Thermo Fisher) and human NEK7 for His-SUMO fusion protein expression in *E. coli* BL21 (DE3) were described recently (Sharif, Wang et al. 2019). For NLRP3 expression the baculovirus of NLRP3 was prepared using the Bac-to-Bac system (Thermo Fisher). Protein expression was induced by infection of Sf9 cells with 1% v/v of baculovirus. 48 h after infection, cells were lysed by sonication in buffer containing 30 mM HEPES, 200 mM NaCl, 2 mM 2-mercaptoethanol and 10% glycerol at pH 7.5 with freshly added protease inhibitor cocktail (Sigma). The supernatant was incubated with 3 ml amylose resin at 4 °C for 1 h and subjected to gravity flow. NLRP3 protein was eluted with 50 mM maltose and further purified with size-exclusion chromatography on Superose 6 10/300 GL column (GE Healthcare) equilibrated with buffer containing 30 mM HEPES, 150 mM NaCl and 2 mM β-mercaptoethanol at pH 7.5. NEK7 was overexpressed in *E. coli* BL21 (DE3) overnight at 18 °C after induction with 0.1 mM isopropyl-β-d-thio-galacto-pyranoside after optical density at 600 nm reached 0.8. Cells were lysed by sonication in buffer containing 50 mM HEPES, 500 mM NaCl, 5 mM $MgCl_2$, 10 mM imidazole, 10% glycerol and 2 mM β-mercaptoethanol at pH 7.5 with freshly added protease inhibitor cocktail (Sigma). The His-SUMO-fusion NEK7 was purified by affinity chromatography using Ni-NTA beads (Qiagen), followed by size-exclusion chromatography on Superdex 200 10/300 GL column (GE Healthcare), equilibrated with buffer containing 30 mM HEPES, 150 mM NaCl and 2 mM β-mercaptoethanol at pH 7.5. WT and kinase-dead mutant BTK were overexpressed in Expi293 cells (Thermo Fisher) using transient transfection with poly-ethylenimine 25K (Polysciences). Cells were harvested 96 h post transfection and lysed in buffer containing 50 mM HEPES, 150 mM NaCl, 2 mM 2-mercaptoethanol and 10% glycerol at pH 7.5 with freshly added protease inhibitor cocktail (Sigma). The FLAG-fusion proteins were subjected to affinity chromatography using anti-FLAG M2 affinity gel (Millipore Sigma), eluted with 3xFLAG-peptide (Millipore Sigma) and further purified by size-exclusion chromatography on Superdex 200 10/300 GL column (GE Healthcare) equilibrated with buffer containing 30 mM HEPES, 150 mM NaCl and 2 mM β-mercaptoethanol at pH 7.5. Proteins were concentrated to 2-7 mg/ml, flash-frozen in liquid nitrogen and stored at -80 °C.

**In vitro pull-downs.** MBP-tagged NLRP3 (2 µM) was mixed with 4 µM His-SUMO-NEK7 or wild type or mutant FLAG-BTK in buffer containing 30 mM HEPES, 150 mM NaCl and 2 mM β-mercaptoethanol at pH 7.5, and incubated for 30 min at 30 °C. The mixture was further incubated for 1 h with 40 µl amylose resin and washed twice with 500 µl of the same buffer, followed by 1 h elution with 50 mM maltose. 30% and 70% of the sample was loaded as input and elution fractions, respectively, and analyzed by SDS–PAGE and immunoblot using monoclonal Anti-Flag® M2-Peroxidase (HRP) or anti-p-Y antibody (Sigma-Aldrich).

**ELISA.** Human and murine IL-1ß, IL-6 or TNF in supernatants were determined by ELISA using half-area plates and kits by R&D Systems and Biolegend, determining triplicate measurements on a standard plate reader.

**Co-immunoprecipitation and immunoblot.** PBMCs or BMDMs were primed with LPS and stimulated with nigericin, washed with cold PBS and immediately lysed in RIPA lysis buffer containing protease/phosphatase inhibitors (Roche). A sample of the cleared lysate was taken before addition of the primary antibody (see Table S2). After 18 hours of rotation at 4 °C, magnetic bead coupled secondary antibody (Protein G Dynabeads, Thermo Fisher) was added for another 90 min. The beads were then washed three times with lysis buffer, resuspended in SDS loading buffer and boiled. HEK293T were transfected using $CaPO_4$ and 24 h later treated with 1 µM MCC950, 60 µM ibrutinib or 60 µM acalabrutinib for 6 h in cases where indicated. Cells were lysed 48 h post-transfection in RIPA buffer supplemented with protease/phosphatase inhibitors (Roche). Cleared lysates were subjected to immunoprecipitation of the NLRP3-HA or NACHT-FLAG fusion protein with Dynabeads (Sigma-Aldrich), or with agarose beads covered with PI4P (P-B004a, Echelon Biosciences). Washed beads were boiled in loading buffer and applied to standard SDS-PAGE on Thermo Fisher pre-cast gels, followed by immunoblot according to the antibody manufacturer's instructions. Membranes were exposed using Peqlab Fusion FL camera and FusionCapt Advance software. Quantification was conducted using the same software.

**WES capillary electrophoresis.** 3 µl of the prepared Immunoblot lysates were run on a ProteinSimple WES instrument, according to the manufacturer´s instructions. Data were analyzed with the Compass for SW software comparing the p-NLRP3 signal with the heavy chain signal from the same run as an internal control.

**Native PAGE.** BMDMs were stimulated and lysed in RIPA lysis buffer without SDS: Lysates were centrifuged at 2,300 x *g* for 10 min to pellet DNA. Supernatant was centrifuged at 16,100 x *g* for 25 min and the pellet was resuspended in native PAGE sample buffer (Thermo Fisher). The samples were loaded onto NuPage 3-8% Tris-Acetate gels (Thermo Fisher) without boiling and native PAGE was conducted using Tris-Glycine running buffer (Thermo Fisher). The gel was soaked in 10% SDS solution for 10 min before performing semi-dry transfer and continuing with conventional immunoblot.

**Crosslinking of ASC oligomers.** BMDMs were primed with LPS and treated with ibrutinib and nigericin. Cells were lysed in RIPA lysis buffer and pellets were cross-linked using DSS and analyzed as described in (Khare, Radian et al. 2016).

**Size exclusion chromatography.** BMDMs were stimulated and lysed in 50 mM Tris-HCl pH 7.4, 1% NP-40, and 150 mM NaCl. 100 µl cleared lysate were loaded on a Superdex 200 Increase 10/300 GL (GE Healthcare) column and proteins were eluted using ÄKTA Purifier (GE Healthcare) and buffer 50 mM Tris-HCL pH 7.4 and 150 mM NaCl with 0.25 ml/min flow. 200 µl fractions were collected and analyzed via Western-Blot.

**In vitro kinase assay.** For results in Fig. 1H, recombinant NLRP3 from Novus Biologicals **(**H00114548-P01) and BTK from Sino Biological (10578-H08B) or Abcam (ab205800) were incubated at 30 °C for 3 h using CST kinase buffer (#9802) in the presence of 2 mM ATP. As a negative control, recombinant Posi-Tag Epitope Tag Protein (Biolegend) was used. Before and after kinase assay samples were boiled and analyzed via SDS PAGE and Western Blot. For results in Fig. S1B, NLRP3 and BTK (WT or KD) were purified as described above. For reactions 2 µM MBP-tagged NLRP3 was mixed with or without 0.2 µM purified FLAG-tagged BTK in buffer containing 30 mM HEPES, 150 mM NaCl, 12.5 mM $MgCl_2$, 2.5 mM ATP and 2 mM β-mercaptoethanol at pH 7.5 in presence or absence of ibrutinib (Selleckchem, cat. S2680). The mixture was incubated at 30 °C and equal aliquots were taken at indicated time points. Samples were analyzed by SDS–PAGE and immunoblot using anti-p-Y antibody (Cell signaling, cat. 8954S).

**Dot blot analysis.** Synthetized peptides were incubated with recombinant BTK (Sino Biologicals) for 3 h in CST kinase buffer (#9802) supplemented with 2 mM ATP. Next, the samples were boiled and anti-His magnetic beads (Dynabeads™ His-Tag Isolation and Pulldown, Thermo Fisher) were added to deplete the samples of phosphorylated BTK. The samples were cleared from the magnetic beads and the supernatants were manually spotted on a nitrocellulose membrane. The dried spots were stained using the Pierce reversible protein stain to visualize total peptide amounts. Then the membrane was blocked with 5% BSA in TBS-T and conventional anti-phospho-Tyrosine primary and secondary antibody incubation steps followed.

**Subcellular fractionation.** Cells were homogenized using a 10 ml syringe and 27 G x 19 mm needles in homogenization buffer (0.25 M sucrose, 10 mM Tris HCl (pH 7.5), 10 mM KCl, 1.5 mM $MgCl_2$, protease inhibitor (Roche) and PhosStop (Roche)). Homogenized cells were centrifuged at 1,000 x *g* for 5 min to remove the nucleus. The supernatant was centrifuged at 5,000 x *g* for 10 min to obtain a heavy membrane fraction (pellet, P5). The supernatant was centrifuged 100,000 x *g* for 20 min to separate a light membrane fraction (S100) from the cytosol. P5 and S100 were washed once with homogenization buffer and then used for sucrose gradient ultracentrifugation, separately. For sucrose gradient ultracentrifugation, a continuous 15-45% (w/w) sucrose gradient was prepared in 10 mM Tris-HCl (pH 7.5), 20 mM KCl, and 3 mM $MgCl_2$) using a Biocomp Gradient Station (Biocomp Instruments). P5 or S100 was loaded on top of the gradient and centrifuged at 170,000 x *g* for 3 h. The gradient was fractionated into 12 fractions of 1.1 ml using the fraction collector module of a Biocomp Gradient Station.

**PI4P bead binding assays**. HEK293T cells were transfected with HA-tagged human WT or Y>E PYD/NACHT linker (residues 94-219)-mCitrine-HA constructs. Cells were lysed in RIPA buffer and PI4P (Echelon Biosciences, P-B004A) or the same amount of control beads (Echelon Biosciences, P-B000) were added to cleared lysates and incubated for 1.5 h at 4 °C while rotating. Beads were then washed 3 times with RIPA buffer, boiled and bound proteins were analyzed via immunoblot. Alternatively, cells were transfected with WT, Y>E or K>A Flag-tagged murine polybasic region (residues 127-146)-GFP-Flag constructs, adopted from (Chen and Chen 2018). PI4P beads or control beads were blocked beforehand in 2% BSA, 0.5% NP-40 and 200 µg/ml Flag peptide (Sigma-Aldrich, F3290) for 2 h at 4°C. Transfected cells were then lysed in RIPA buffer and the expressed proteins were purified using Anti-FLAG® M2 Magnetic Beads (M8823, Merck). Beads were washed 3x with RIPA buffer and boiled to elute the purified polybasic region. Blocked PI4P beads or the same amount of control beads were added to the eluted protein, and incubated for 1.5 h on 4 °C while rotating. Beads were then washed 3 x 8 min with RIPA buffer, resuspended in LDS sample buffer, boiled, and bound protein was analyzed using immunoblot.

**Reconstitution and analysis of NLRP3-deficient immortalized macrophages.** NLRP3-deficient immortalized macrophages (Hornung, Bauernfeind et al. 2008) were retrovirally transduced with NLRP3 (WT or 4xY>F)-Flag-T2A-mCherry constructs as described in (Hornung, Bauernfeind et al. 2008) and subsequently sorted for similar expression of mCherry. Similar NLRP3 expression was confirmed by anti-Flag immunoblot of cell lysates. Cells were seeded at $1x10^5$ cells/well in a 96 well plate in 100 µl, primed with LPS (200 ng/ml) for 3h and inflammasome stimuli in optiMEM added as follows: nigericin at 8 µM for 1.5 h, R837 (imiquimod) at 20 µg/ml for 2 h or poly(dA:dT) at 200 ng per well with 0.5 µl lipofectamine 2000 for 4 h. IL-1β and TNF were subsequently determined by IL-1β and TNF HTRF assay respectively (Cisbio; 62MIL1BPEH and 62MTNFAPEG).

**NLRP3 sequence analysis, structure inspection and charge prediction.** NLR sequences were retrieved from UniProt and ClustalW aligned within Geneious R6 software. A hypothetical active conformation of NLRP3 was modeled based on NLRP3-NEK7 structure in an inactive state (PDB 6NPY) (Sharif, Wang et al. 2019). NACHT domain reorganization and hypothetical NLRP3 oligomerization was generated, based on the NLRC4 oligomer (PDB 3JBL) as a homology model template by introduction of a 90° rotation of NBD-HD1 module (Sharif, Wang et al. 2019). Phosphorylation of tyrosine residues of interest was performed in Pymol (Schrödinger) using the PyTMs plugin *(Warnecke, Sandalova et al. 2014)*. Electrostatic potential of the solvent accessible surface of phosphorylated and non-phosphorylated NLRP3 models was calculated with PBEQ-Solver online visualization tool (http://www.charmm-gui.org (MacKerell 1998; Jo, Kim et al. 2008; Jo, Vargyas et al. 2008) and visualized with Pymol. Protein net charges of the Y136, Y140 and Y143-containing linker were conducted with ProtPi (www.protpi.ch).

**Statistics.** Experimental data was analyzed using Excel 2010 (Microsoft) and/or GraphPad Prism 6, 7 or 8, microscopy data with ImageJ/Fiji, flow cytometry data with FlowJo 10. Normal distribution in each group was always tested using the Shapiro-Wilk test first for the subsequent choice of a parametric (ANOVA, Student's *t*-test) or non-parametric (e.g. Friedman, Mann-Whitney U or Wilcoxon) test. p-values ($\alpha$=0.05) were then calculated and multiple testing was corrected for in Prism, as indicated in the figure legends. Values <0.05 were generally considered as statistically significant and denoted by * throughout. Comparisons were made to unstimulated control, unless indicated otherwise, denoted by brackets.

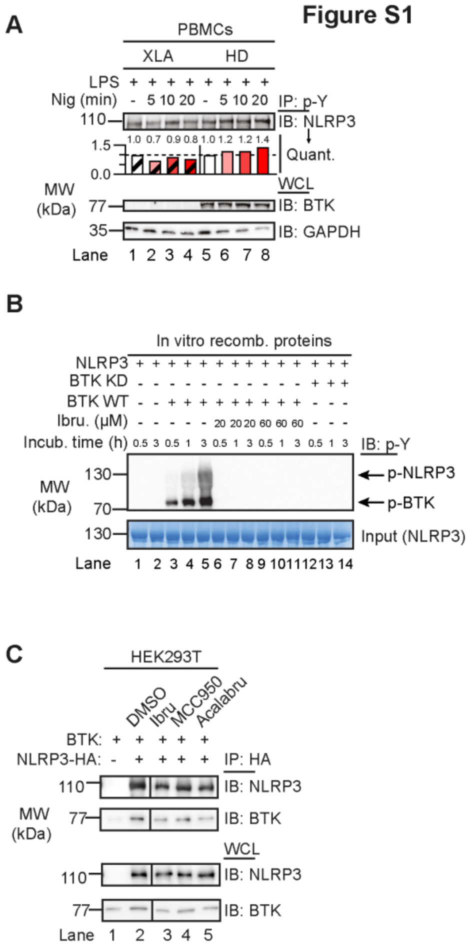


**Figure S1**: (A) Co-IP of NLRP3 from healthy donor (HD) or XLA patients´ PBMC lysates (n=3). (B) p-NLRP3 occurrence in the *in vitro* kinase assay with BTK or kinase-dead BTK (KD) upon incubation with ATP for the indicated periods of time, with and without ibrutinib. (C) HEK293T cells were transfected with the indicated NLRP3 and BTK WT or mutant constructs, treated with inhibitors and lysates subjected to HA-IP and immunoblot (n=3 each). In A-C one representative of ‘n’ biological replicates is shown.

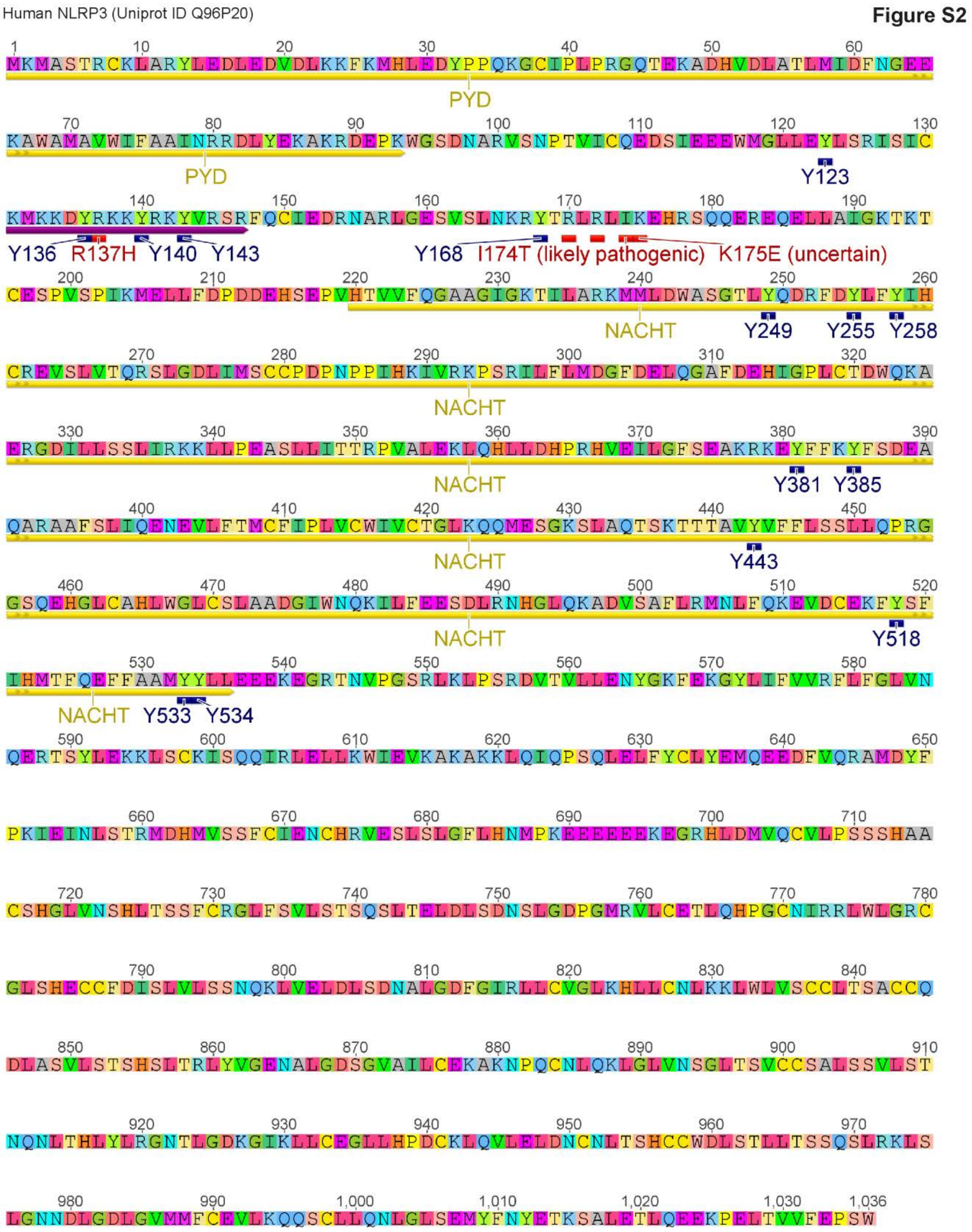


**Figure S2:** Annotation of NLRP3 sequence (UniProt ID Q96P20).

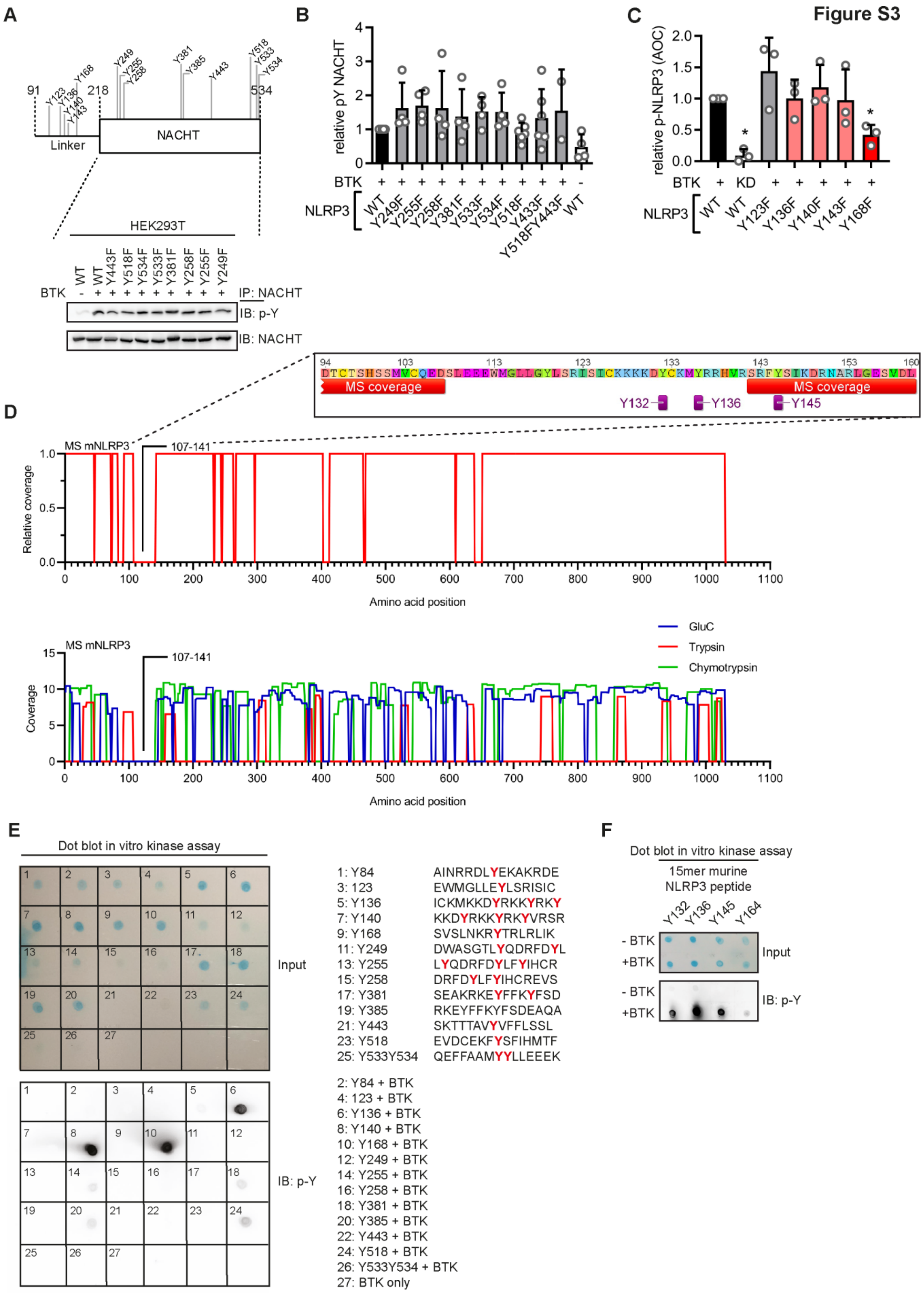
Figure S3
A
B
C
D
E
F
Linker
NACHT
HEK293T
IP: NACHT
IB: p-Y
IB: NACHT
relative pY NACHT
relative p-NLRP3 (AOC)
BTK
NLRP3
MS coverage
MS mNLRP3
107-141
Relative coverage
Coverage
Amino acid position
GluC
Trypsin
Chymotrypsin
Dot blot in vitro kinase assay
Input
IB: p-Y
1: Y84 AINRRDLYEKAKRDE
3: 123 EWMGLLEYLSRISIC
5: Y136 ICKMKKDYRKKYRKY
7: Y140 KKDYRKKYRKYVRSR
9: Y168 SVSLNKRYTRLRLIK
11: Y249 DWASGTLYQDRFDYL
13: Y255 LYQDRFDYLFYIHCR
15: Y258 DRFDYLFYIHCREVS
17: Y381 SEAKRKEYFFKYFSD
19: Y385 RKEYFFKYFSDEAQA
21: Y443 SKTTTAVYVFFLSSL
23: Y518 EVDCEKFYSFIHMTF
25: Y533Y534 QEFFAAMYYLLEEEK
2: Y84 + BTK
4: 123 + BTK
6: Y136 + BTK
8: Y140 + BTK
10: Y168 + BTK
12: Y249 + BTK
14: Y255 + BTK
16: Y258 + BTK
18: Y381 + BTK
20: Y385 + BTK
22: Y443 + BTK
24: Y518 + BTK
26: Y533Y534 + BTK
27: BTK only
15mer murine NLRP3 peptide
Y132
Y136
Y145
Y164
- BTK
+BTK

**Figure S3:** (A) Position of all mutated tyrosine residues in Linker-NACHT construct. (B) Phosphorylation analysis of core-NACHT tyrosine NLRP3 mutants. HEK293T cells were transfected with the indicated NLRP3 mutant constructs and a BTK WT construct as indicated and lysates subjected to HA-IP and immunoblot as indicated (n=4 each). (B) Quantification of A combined from n=4 experiments. (C) Quantification of WES capillary electrophoresis of NLRP3 p-Y IPs from HEK293T cells (Fig. 2H) from n=3 experiments. (D) Mass spectrometric analysis of purified mNLRP3, digested with different proteases, showing combined (upper) and separate coverage (lower) information (Ref. (Stutz, Horvath et al. 2013)). (E) Dot blot of *in vitro* kinase assay of His-BTK and 15-mer synthetic peptides derived from human NLRP3, containing the indicated tyrosines stained with a total protein stain (upper; input) or anti-p-Y Abs (lower grid, n=3). (F) As in E but with peptides derived from murine NLRP3 (n=3). B, D represent combined data (mean+SD) from 'n' biological replicates. In E-F one representative of 'n' biological replicates is shown. * $p<0.05$ according to one sample *t*-test (B, C).

Figure S4

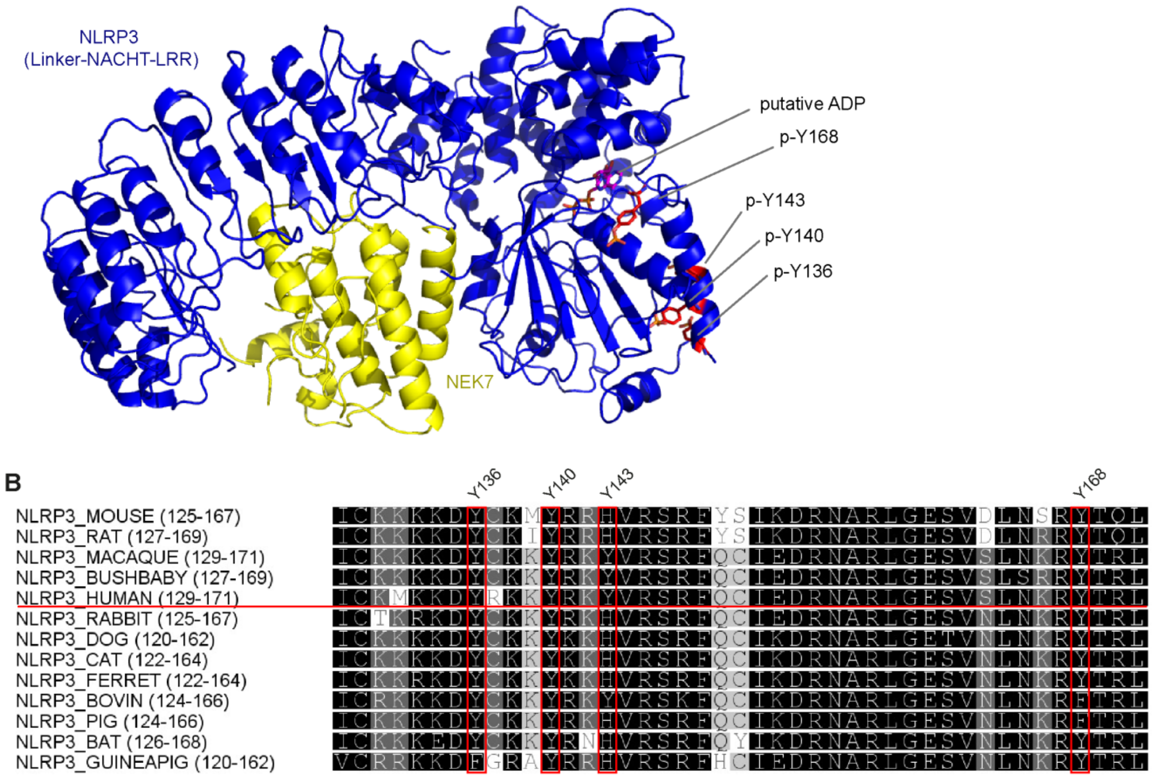


**Figure S4:** Structural aspects and conservation of BTK-modified tyrosine residues in NLRP3. (A) NLRP3 model 6NPY showing NLRP3 Linker-NACHT-LRR (blue) and NEK7 C-terminal lobe (yellow). A putative bound ADP molecule and selected tyrosines are highlighted. (B) ClustalW multiple sequence alignments of NLRP3 sequences from other species. Coloring according to similarity (black = conserved). BTK-modified tyrosines are highlighted (residue numbering according to human NLRP3).

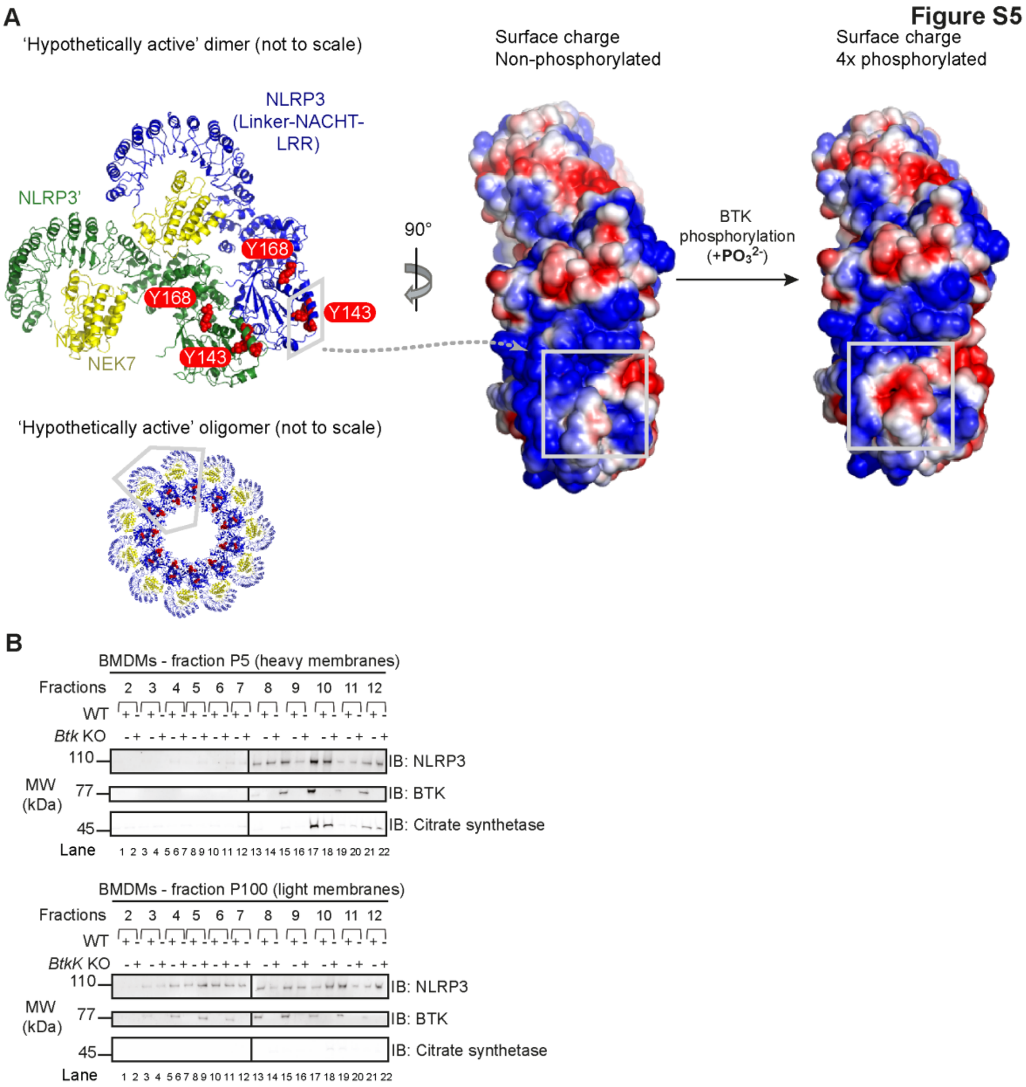


**Figure S5:** (A) CHARMM surface charge predictions of Linker-NACHT-LRR structure in the non-phosphorylated (left) and 4x phosphorylated (right) form. Blue = positive, red = negative charge. Grey boxes indicate that the area of charge alterations in the monomers maps to a contact area in the hypothetical dimer (center, rotated by 90 °; see relative position in oligomer, below). (B) Sucrose gradient fractionation of WT and Btk KO BMDM lysates upon LPS + nigericin treatment for 5 min. Fractions were analyzed by SDS-PAGE and immunoblot as indicated (n=1; pilot fractionation experiment).

Figure S6

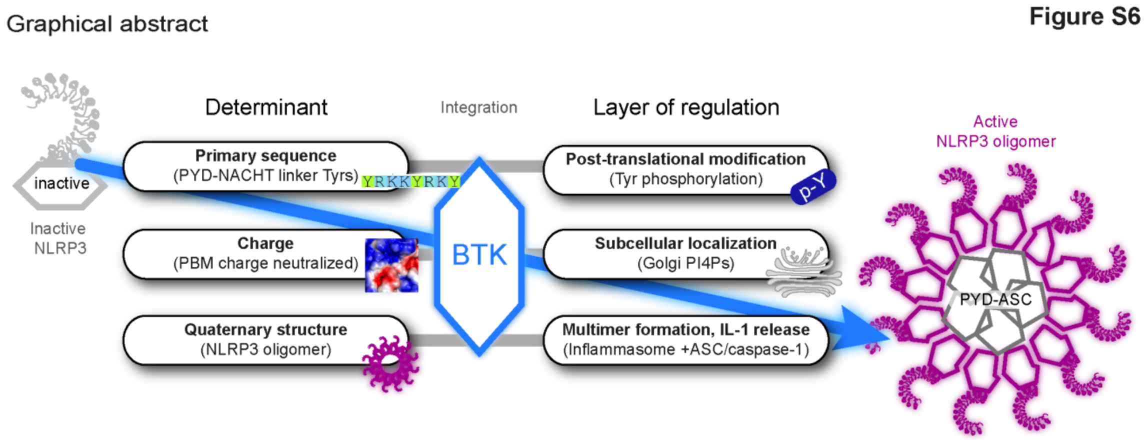


**Figure S6:** Graphical summary illustrating multiple roles of BTK as NLRP3 regulator.

**Table S1:** Peptides used in this study.

| No. | Contained tyrosine(s) | Sequence | Modifications | Use in Fig. | Manufacturer |
|---|---|---|---|---|---|
| | | Human NLRP3 | | | |
| 1 | Y84 | AINRRDLYEKAKRDE | | S3 | In house |
| 2 | Y123 | EWMGLLEYLSRISIC | | S3 | In house |
| 3 | Y136, Y140, Y143 | ICKMKKDYRKKYRKY | | S3 | In house |
| 4 | Y136, Y140, Y143 | KKDYRKKYRKYVRSR | | F2, F3 | In house |
| 5 | Y136, Y140, Y143 | YRKKYRKYVRSRFQC | | S3 | In house |
| 6 | Y136 | KKDYRKKFRKFVRSR | Y140F, Y143F | F2 | In house |
| 7 | Y140 | ICKMKKDFRKKYRKF | Y136F, Y143F | F2 | In house |
| 8 | Y143 | FRKKFRKYVRSRFQC | Y136F, Y140F | F2 | In house |
| 9 | Y168 | SVSLNKRYTRLRLIK | | F3, S3 | In house |
| 10 | Y249 | DWASGTLYQDRFDYL | | S3 | In house |
| 11 | Y255 | LYQDRFDYLFYIHCR | | S3 | In house |
| 12 | Y258 | DRFDYLFYIHCREVS | | S3 | In house |
| 13 | Y381 | SEAKRKEYFFKYFSD | | S3 | In house |
| 14 | Y385 | RKEYFFKYFSDEAQA | | S3 | In house |
| 15 | Y443 | SKTTTAVYVFFLSSL | | S3 | In house |
| 16 | Y518 | EVDCEKFYSFIHMTF | | S3 | In house |
| 17 | Y533, Y534 | QEFFAAMYYLLEEEK | | S3 | In house |
| 18 | Y136, Y140, Y143 | KKDFRKKFRKFVRSR | Y136F, Y140F, Y143F | F2 | In house |
| 19 | Y168 | SVSLNKRFTRLRLIK | Y168F | F2 | In house |
| 20 | Y136, Y140, Y143 | KKDpYRKKpYRKpYVRSR | Phospho-Y | F3 | In house |
| | | Murine NLRP3 | | | |
| 21 | Y132 | ICKKKKDYCKMFRRH | Y136F | S3 | In house |
| 22 | Y136 | KKDFCKMYRRHVRSR | Y132F | S3 | In house |
| 23 | Y144 | RHVRSRFYSIKDRNA | | S3 | In house |
| 24 | Y164 | SVDLNSRYTQLQLVK | | S3 | In house |
| 25 | Y132, Y136 | ICKKKKDYCKMYRRH | | F3 | In house |
| 26 | Y132, Y136 | ICKKKKDpYCKMpYRRH | Phospho-Y | F3 | In house |

**Table S2:** Antibodies used in this study.

| Specificity | Manufacturer | Cat. No. |
|---|---|---|
| mNLRP3 | Adipogen | AG-20B-0014-C100 |
| hNLRP3 | CST | #15101 |
| hBTK | BD | 611117 |
| mBTK | CST | #8547 |
| HA | CST | #3724 |
| Flag | Sigma-Aldrich | F1804 |
| p-Y | CST | #8954 |
| ASC | CST | #67824 |
| hIL-1β | R&D Systems | MAB601 |
| mIL-1β | CST | #12242 |
| mCaspase-1 | CST | #3866 |
| GAPDH | Thermo-Fisher | # MA5-15738 |
| Citrate synthetase | GeneTex | GTX110624 |
| Anti-Mouse IgG (H+L) HRP conjugate | Promega | W4021 |
| Anti-Rabbit IgG (H+L) HRP conjugate | Vector | PI-1000 |
| VeriBlot for IP Detection Reagent (HRP) | Abcam | Ab131366 |
| Anti-Flag® M2-Peroxidase (HRP) | Sigma-Aldrich | A8592 |
